# LUBAC ubiquitinates FADD and Caspase-8 through distinct catalytic mechanisms and limits TNF-induced apoptosis

**DOI:** 10.64898/2026.11.22.768392

**Authors:** Frederick Kumi-Ansah, Arianna Borrego, Jaime Lopez-Mosqueda

## Abstract

The linear ubiquitin chain assembly complex (LUBAC) is a critical regulator of tumor necrosis factor (TNF) signaling that promotes pro-survival responses and suppresses apoptosis. Linear ubiquitination of the TNFR1-associated complex II components FADD and Caspase-8 has been reported, but whether they are direct LUBAC substrates, how they are modified, and what the modification does remain poorly understood. Here, we show that loss of HOIP, which destabilizes the LUBAC complex, sensitizes HCT116 cells to TNF-induced apoptosis, and that TNF induces linear ubiquitin-reactive species on endogenous FADD and Caspase-8. Using purified proteins, we demonstrate that FADD and Caspase-8 are direct LUBAC substrates with different catalytic requirements. Efficient FADD ubiquitination requires HOIL-1L in addition to HOIP, consistent with a priming-and-extension mechanism, whereas Caspase-8 is ubiquitinated by HOIP-containing subcomplexes lacking HOIL-1L. A complex containing FADD, Caspase-8, and RIPK1 formed independently of LUBAC activity, but pharmacological inhibition of HOIP increased Caspase-8 processing within the FADD-associated pool, and Met1-ubiquitinated FADD showed reduced binding to Caspase-8 in vitro. Mapping identified four death domain lysines that each support LUBAC-dependent FADD ubiquitination. In FADD-deficient cells, a FADD-diubiquitin fusion attenuated TNF/cycloheximide-induced apoptosis relative to wild-type FADD, whereas a lysine-less FADD supported apoptosis with delayed kinetics. Together, these findings identify FADD and Caspase-8 as direct LUBAC substrates modified through distinct catalytic routes and support a model in which ubiquitination of FADD limits productive engagement with Caspase-8.

## Introduction

Tumor necrosis factor (TNF) is a pleiotropic cytokine that regulates inflammation, immunity, cell survival, and cell death^1^. Binding of TNF to the TNF receptor 1 (TNFR1) triggers the assembly of two distinct signaling complexes^2^. At the membrane, TRADD, TRAF2, RIPK1, and the E3 ubiquitin ligases cIAP1/2 assemble into complex I, which promotes NF-κB activation and pro-survival gene expression^2–5^. When complex I signaling is compromised, its components can instead nucleate a cytosolic complex (complex II), in which RIPK1 and FADD recruit and activate Caspase-8 to drive apoptosis^2,6,7^. The balance between these outcomes is controlled by post-translational modification, particularly ubiquitination, which governs the assembly, stability, and signaling output of both complexes^8–10^.

The linear ubiquitin chain assembly complex (LUBAC) is a central regulator of complex I signaling^6^. LUBAC comprises the catalytic E3 ligase HOIP together with the subunits HOIL-1L and SHARPIN^11–14^. LUBAC-generated linear chains on NEMO and RIPK1 stabilize complex I and are required for optimal NF-κB activation^15^, and LUBAC loss also sensitizes cells to TNF-induced apoptosis^12–14^. Underscoring the physiological importance of this pathway, LUBAC-deficient mice die at mid-gestation from excess endothelial cell death^16^, and humans with inherited HOIL-1L deficiency develop severe autoinflammation and immunodeficiency^17^.

Much of LUBAC’s anti-apoptotic function has been attributed to its role in stabilizing complex I and sustaining pro-survival gene expression. However, RIPK1, FADD, and Caspase-8 have each been reported to acquire linear ubiquitin modifications following TNF or TRAIL stimulation^18–21^, raising the possibility that LUBAC also acts directly on complex II itself. These studies, however, have relied largely on ubiquitination detected in cells, and have not established which of these proteins are direct LUBAC substrates, through what catalytic mechanism they are modified, or what functional consequence this modification has for complex II assembly and signaling. Whether LUBAC restrains apoptosis solely by limiting complex II formation upstream, or additionally by directly modifying complex II components once formed, therefore remains an open question.

FADD serves as the central adaptor of complex II, tethering RIPK1 and death receptors to procaspase-8 through homotypic death domain and death effector domain interactions^2,22,23^. The recruitment and dimerization of procaspase-8 at this platform triggers its autoproteolytic activation and initiates the apoptotic caspase cascade^24^. Because FADD and Caspase-8 occupy this pivotal position, any post-translational modification that alters their conformation, interactions, or availability has the potential to determine whether a cell survives or dies in response to TNF. Whether linear ubiquitination directly regulates assembly or signaling competence of the FADD–Caspase-8 module has not, to our knowledge, been directly tested.

Recent work has shown that HOIL-1L and HOIP can play distinct catalytic roles during substrate modification^25,26^. HOIL-1L conjugates an initial monoubiquitin onto a substrate, which HOIP then extends into a linear chain, a priming-and-extension mechanism described for LUBAC’s own subunits and for NEMO^27,28^. Whether this two-step process applies uniformly across LUBAC substrates, or whether individual substrates are recognized and modified through distinct catalytic routes, is unknown. Resolving this question is important for understanding how a single ligase complex achieves selective, substrate-specific control over multiple, mechanistically distinct branches of TNF signaling.

In the present study, we investigated how LUBAC regulates TNF-induced apoptotic signaling. We show that HOIP loss sensitizes HCT116 cells to TNF-induced apoptosis and that TNF induces linear ubiquitin-reactive species on endogenous FADD and Caspase-8. Using purified components, we show that FADD and Caspase-8 are both direct substrates of LUBAC but have different catalytic requirements. FADD ubiquitination requires the coordinated, sequential activities of HOIL-1L and HOIP, consistent with a priming-and-extension mechanism, whereas Caspase-8 is ubiquitinated by HOIP-containing subcomplexes that lack HOIL-1L. We further show that ubiquitination of FADD reduces its binding to Caspase-8 in vitro, and that a FADD-ubiquitin fusion attenuates apoptotic signaling. Together, these findings identify substrate-specific catalytic steps in LUBAC-mediated ubiquitination of complex II components and suggest that LUBAC may establish a ubiquitin-dependent checkpoint on complex II.

## Results

### HOIP loss sensitizes cells to TNF-induced apoptosis, and TNF induces linear ubiquitination of FADD and Caspase-8

To establish the physiological context for this study, we first generated HOIP-deficient HCT116 cells using CRISPR/Cas9 (**Figure 1A**). Loss of HOIP was accompanied by a reduction in HOIL-1L and SHARPIN protein levels (**Figure S1A**), consistent with the known requirement for an intact heterotrimeric complex to maintain expression of the individual LUBAC subunits^12–14^.

**Figure 1.**
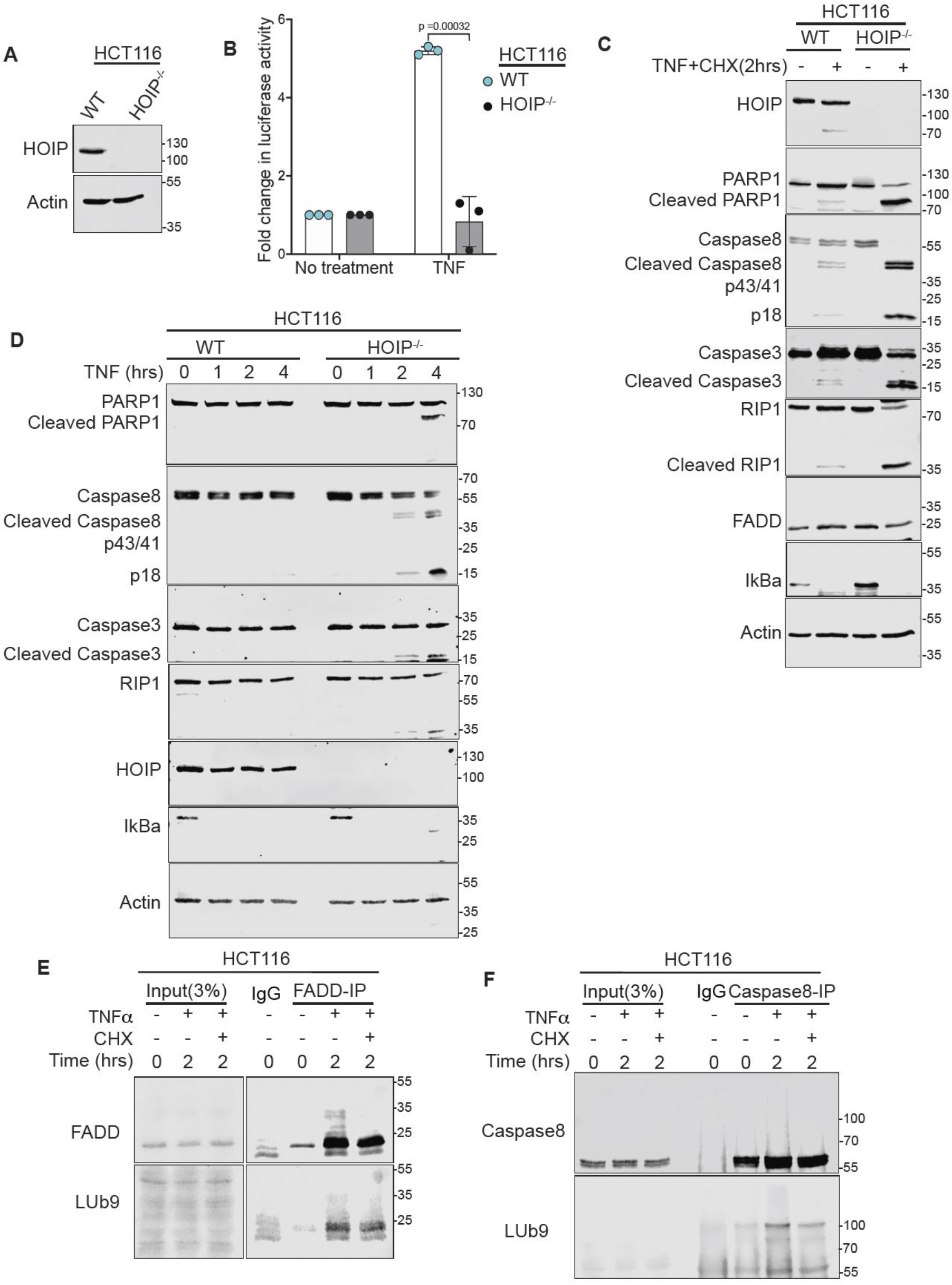
HOIP loss sensitizes cells to TNF-induced apoptosis, and TNF induces linear ubiquitin-reactive species on FADD and Caspase-8. **A)** Immunoblot confirming loss of HOIP in CRISPR/Cas9-generated HOIP-KO HCT116 cells. Actin, loading control. **B)** NF-κB luciferase reporter activity in WT and HOIP-KO HCT116 cells left untreated or treated with TNF [20 ng/mL, 12 h]. Data are mean ± SD of n = 3; p value by Multiple unpaired t-test with statistical significance determined using Benjamini-Hochberg False Discovery Rate (FDR). **(C)** Immunoblots of WT and HOIP^-/-^HCT116 cells treated with TNF [20 ng/mL] plus cycloheximide [20 µg/mL] for 2 h, probing apoptotic markers (PARP1, caspase-8, caspase-3, RIP1 cleavage), FADD and IκBα. **(D)** Immunoblots of WT and HOIP-KO cells treated with TNF alone [20 ng/mL] for 0-4 h, probed as in C. **(E, F)** Denaturing immunoprecipitation of FADD (E) or Caspase-8 (F) from HCT116 cells treated with TNF [20 ng/mL] ± CHX [20 µg/mL] for 2 h, immunoblotted with a linear ubiquitin-specific antibody (LUb9). Input, 3%; IgG, control IP. Blots are representative of at least two independent experiments.

To confirm that this loss of LUBAC expression translates into a functional defect in canonical LUBAC signaling, we monitored TNF-induced NF-κB activity using a luciferase reporter (**Figure 1B**). TNF stimulation induced reporter activity in wild-type cells not in HOIP-knockout (KO) cells, (∼5.2-fold vs. ∼0.8-fold (p = 0.00032), consistent with the established role of LUBAC in canonical TNF-induced NF-κB signaling^11,15^. HOIP-KO cells therefore lack both LUBAC integrity and signaling output.

We next examined the consequences of LUBAC loss for TNF-induced apoptosis, using two complementary stimulation paradigms. First, we treated cells with TNF in combination with cycloheximide (CHX), a protein-synthesis inhibitor commonly used to unmask apoptotic signaling by preventing synthesis of pro-survival factors^29,30^. Under these acute conditions (2 hours), HOIP-KO cells showed markedly enhanced cleavage of PARP1, Caspase-8 (generating the p43/41 fragments) and RIPK1, relative to wild-type cells treated in parallel (**Figure 1C**). Second, to determine whether LUBAC loss sensitizes cells to apoptosis even without pharmacological suppression of survival gene expression, we treated cells with TNF alone over a longer time course (0-4 hours). Wild-type cells remained largely resistant to TNF alone throughout this period, whereas HOIP-KO cells progressively accumulated processed Caspase-8 (p43/41 and the fully processed p18 fragment), cleaved PARP1, and cleaved Caspase-3, with these changes clearly apparent by 4 hours (**Figure 1D**). In both experiments, TNF stimulation induced degradation of IκBα, confirming effective TNF signaling in wild-type cells despite the absence of apoptotic marker cleavage (**Figure 1C, 1D**). Cycloheximide accelerates a death program LUBAC-deficient cells are already predisposed to execute.

To determine whether these biochemical markers of apoptosis reflect a greater number of cells undergoing apoptosis, we quantified cleaved Caspase-3-positive cells by immunofluorescence across non-treated, TNF-alone and TNF+ cycloheximide (CHX) conditions (**Figure S1B and S1C**). Wild-type cultures remained essentially free of cleaved Caspase-3 staining under all three conditions. In contrast, HOIP-KO cells showed a clear treatment-dependent increase, from ∼0% (untreated) to ∼7% (TNF) to ∼35% (TNF+ CHX), confirming at the single-cell level that loss of LUBAC sensitizes HCT116 cells to TNF-induced apoptotic death, and that this sensitization is substantially amplified when new protein synthesis is blocked.

To ask whether LUBAC acts directly on components of complex II, rather than acting solely upstream to influence its formation, we immunoprecipitated endogenous FADD and Caspase-8 from HCT116 cells under denaturing conditions and probed with a Met1-linkage-specific antibody (LUb9) (**Figure 1E and 1F**). A 2-hour TNF stimulation induced the appearance of higher-molecular-weight, LUb9-reactive species of both FADD and Caspase-8, indicating that both proteins are modified with linear ubiquitin in cells in a TNF-dependent manner. Co-treatment with cycloheximide markedly reduced this signal for Caspase-8, and to a more modest degree for FADD, indicating that accumulation of these modified species is associated with conditions that favor survival signaling rather than with the acute induction of apoptosis. This pattern, induced by TNF and reduced under strongly pro-apoptotic conditions, is compatible with a role for linear ubiquitination of FADD and Caspase-8 in restraining apoptotic signaling, although it could also reflect loss of LUBAC activity during apoptosis, for example through caspase-mediated cleavage of HOIP^21^. Together with the apoptosis phenotypes described above, these results are consistent with a model in which LUBAC restrains TNF-induced apoptosis in HCT116 cells, at least in part, through TNF-dependent linear ubiquitination of FADD and Caspase-8.

### FADD and Caspase-8 are direct LUBAC substrates with distinct requirements for HOIL-1L in vitro

To determine whether FADD and Caspase-8 are direct substrates of LUBAC, we first examined their ability to interact with individual LUBAC components. In GST pulldown assays, GST-FADD efficiently recovered Caspase-8-FLAG, consistent with their established association, but failed to recover detectable amounts of HOIP-FLAG, HOIL-1L-FLAG, or SHARPIN-FLAG (**Figure 2A**). To determine whether ubiquitination influences LUBAC recruitment, we generated monoubiquitinated FADD chimera (GST-FADD-Ub). Unlike unmodified FADD, GST-FADD-Ub recovered HOIP while retaining its interaction with Caspase-8, whereas interactions with SHARPIN and HOIL-1L remained undetectable (**Figure 2A**). These findings suggest that ubiquitin conjugation promotes an interaction between FADD and HOIP.

**Figure 2.**
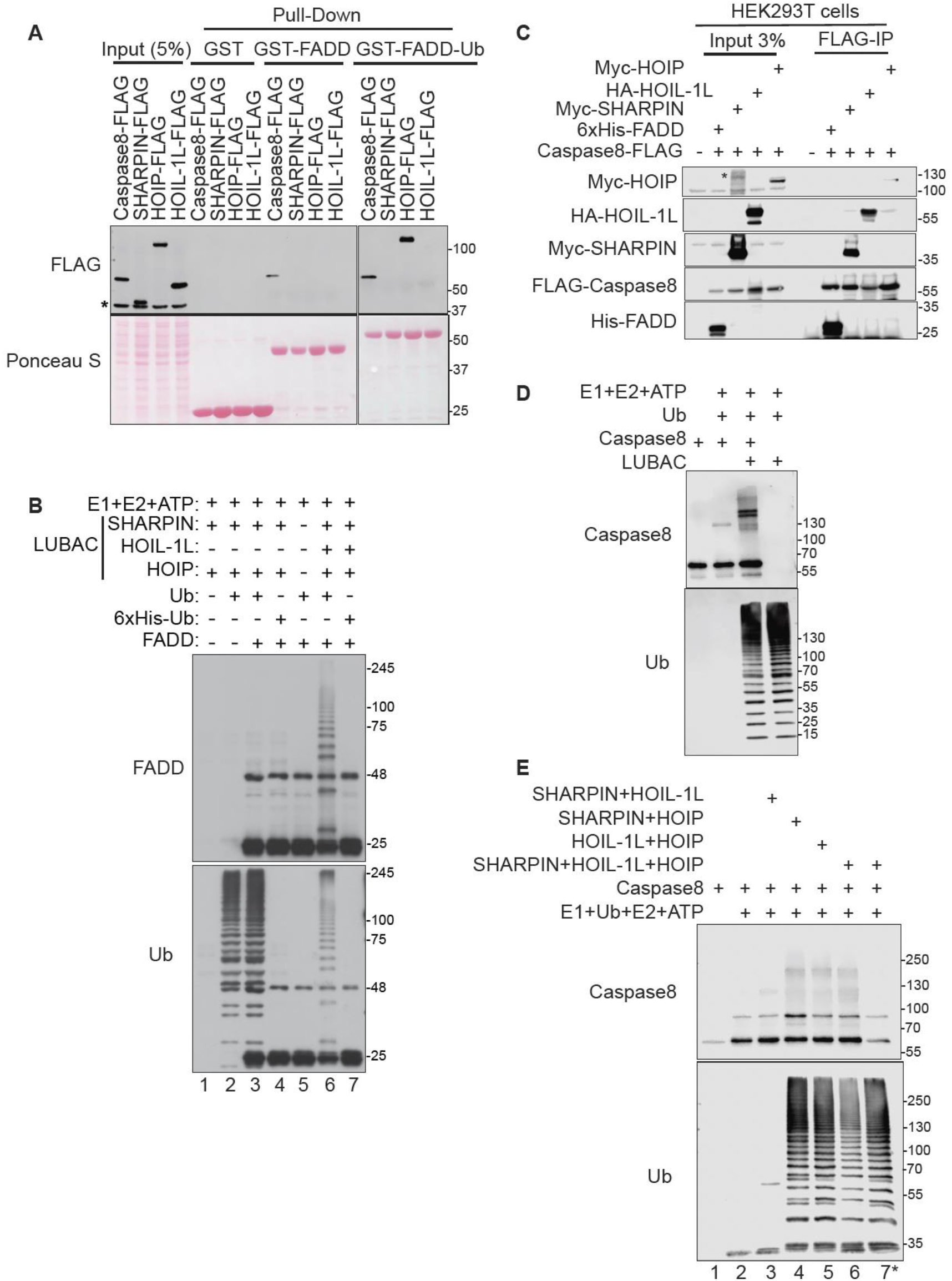
FADD and caspase-8 are LUBAC substrates in vitro. **(A)** GST pull-down of FLAG-tagged Caspase-8, SHARPIN, HOIP or HOIL-1L, transiently expressed in HEK293T cells, using GST (negative control), GST-FADD or GST-FADD-Ub (monoubiquitinated FADD; FADD-Ubiquitin fusion) as bait. Caspase-8 is the positive control for FADD binding. Bound proteins were detected by anti-FLAG immunoblot; Ponceau S shows bait protein. Input, 5%. Asterisk (*), non-specific band. **(B)** In vitro ubiquitination of FADD with E1 (UBE1), E2 (UBCH7), ATP and either ubiquitin (Ub) or N-terminally 6xHis-tagged Ub, which blocks linear ubiquitin chain formation, in the presence of the indicated LUBAC components (lanes 1-7), immunoblotted for FADD and Ub. **(C)** FLAG immunoprecipitation from HEK293T cells co-expressing Caspase-8-FLAG with Myc-HOIP, HA-HOIL-1L, Myc-SHARPIN or 6xHis-FADD (positive control), as indicated. Input, 3%. **(D)** In vitro ubiquitination of Caspase-8 with E1 (UBE1), E2 (UbcH5a), ATP, Ub and LUBAC purified from HEK293T cells, immunoblotted for Caspase-8 and Ub. **(E)** In vitro ubiquitination of Caspase-8 with the indicated LUBAC subcomplexes or full LUBAC (lanes 1-7). In lane 7*, Caspase-8 was added after incubation and the reaction was immediately quenched, to control for Caspase-8 immunoblot signal and for antibody cross-reactivity with ubiquitin chains. Blots are representative of at least two independent experiments.

We next asked whether FADD could be directly modified by LUBAC. To enable fully reconstituted biochemical assays, we expressed and purified full-length HOIP, HOIL-1L, and SHARPIN and assembled them into the heterotrimeric LUBAC complex (**Figure S2A**). In reconstituted ubiquitination reactions containing purified E1, E2, ubiquitin, ATP, and recombinant LUBAC, FADD was converted into a high-molecular-weight laddered species, demonstrating that FADD is a direct LUBAC substrate (**Figure 2B**). To determine the nature of the ubiquitin chains assembled on FADD, ubiquitinated FADD was treated with the Met1-specific deubiquitinase OTULIN. Catalytically active OTULIN efficiently collapsed the high-molecular-weight FADD species, whereas the inactive OTULIN-C129A mutant had no effect, confirming the presence of Met1-linked ubiquitin chains (**Figure S2B**). As an independent validation of OTULIN’s specificity, a panel of deubiquitinase with distinct chain linkage preference (USP2, AMSH, and OTUB1) was applied to preassembled linear ubiquitin chains. OTULIN, but not AMSH or OTUB1, disassembled linear chains, confirming that OTULIN selectively targets linear ubiquitin chains under these conditions (**Figure S2C**). To further validate the reconstituted system, reactions were performed in parallel using NEMO, a well-established LUBAC substrate, or GST alone. NEMO was ubiquitinated under the same conditions that modified FADD, whereas GST remained unmodified, demonstrating both substrate specificity and proper activity of the reconstituted LUBAC complex (**Figure S2D and S2E**).

We next performed analogous experiments with Caspase-8. HEK293T cells were co-transfected with Caspase-8-FLAG together with 6xHIS-FADD, SHARPIN, HOIP, or HOIL-1L individually. Caspase-8-FLAG was immunoprecipitated from each lysate. Caspase-8-FLAG recovered FADD, used here as a positive control, as well as HOIL-1L and SHARPIN, while HOIP was recovered less efficiently (**Figure 2C**). To assess whether ubiquitination influences these interactions, we generated a Caspase-8-Ub chimera and examined its binding to individual LUBAC subunits. Similar to the interaction profile of unmodified Caspase-8, the ubiquitinated chimera associated with SHARPIN and HOIL-1L showed a weaker interaction with HOIP (**Figure 2D**). Thus, unlike FADD, Caspase-8 displays detectable interaction with LUBAC components even in the absence of pre-existing ubiquitin modification.

We next tested whether Caspase-8 is a direct LUBAC substrate. In reconstituted ubiquitination reactions containing the complete LUBAC complex, purified Caspase-8 was efficiently converted into a high-molecular-weight laddered species, demonstrating direct ubiquitination by LUBAC (**Figure 2E**). Having established that both FADD and Caspase-8 are direct LUBAC substrates, we next compared the LUBAC components required for substrate modification. Whereas FADD ubiquitination was strongly dependent on HOIL-1L (**Figure 2B**), Caspase-8 was efficiently ubiquitinated by both HOIP-SHARPIN and HOIP-HOIL-1L subcomplexes, as well as by the complete LUBAC complex (**Figure 2E**). These results indicate that HOIP is sufficient to drive Caspase-8 ubiquitination, whereas efficient FADD ubiquitination requires HOIL-1L. Together, these findings establish FADD and Caspase-8 as LUBAC substrates and reveal distinct mechanisms of substrate modification. While FADD exhibits little interaction with individual LUBAC subunits unless ubiquitinated and requires HOIL-1L for efficient modification, Caspase-8 associates with LUBAC components directly and can be ubiquitinated by HOIP-containing subcomplexes in the absence of HOIL-1L.

### LUBAC activity limits Caspase-8 processing in complex II, and ubiquitination reduces FADD binding to Caspase-8

To determine the functional consequence of FADD ubiquitination, we examined complex II assembly under conditions of LUBAC loss or pharmacological inhibition. We first immunoprecipitated endogenous FADD from wild-type and HOIP-KO cells following TNF and cycloheximide treatment in the presence of the pan-caspase inhibitor Z-VAD-fmk, which stabilizes complex II by blocking Caspase-8-dependent processing and turnover of its components, allowing the assembled complex to be captured^2^. FADD-IP recovered both RIPK1 and Caspase-8 specifically at two hours into TNF/CHX/Z-VAD treatment in both genotypes, confirming that FADD nucleates a RIPK1/Caspase-8-containing complex under these conditions (**Figure 3A and 3B**).

**Figure 3.**
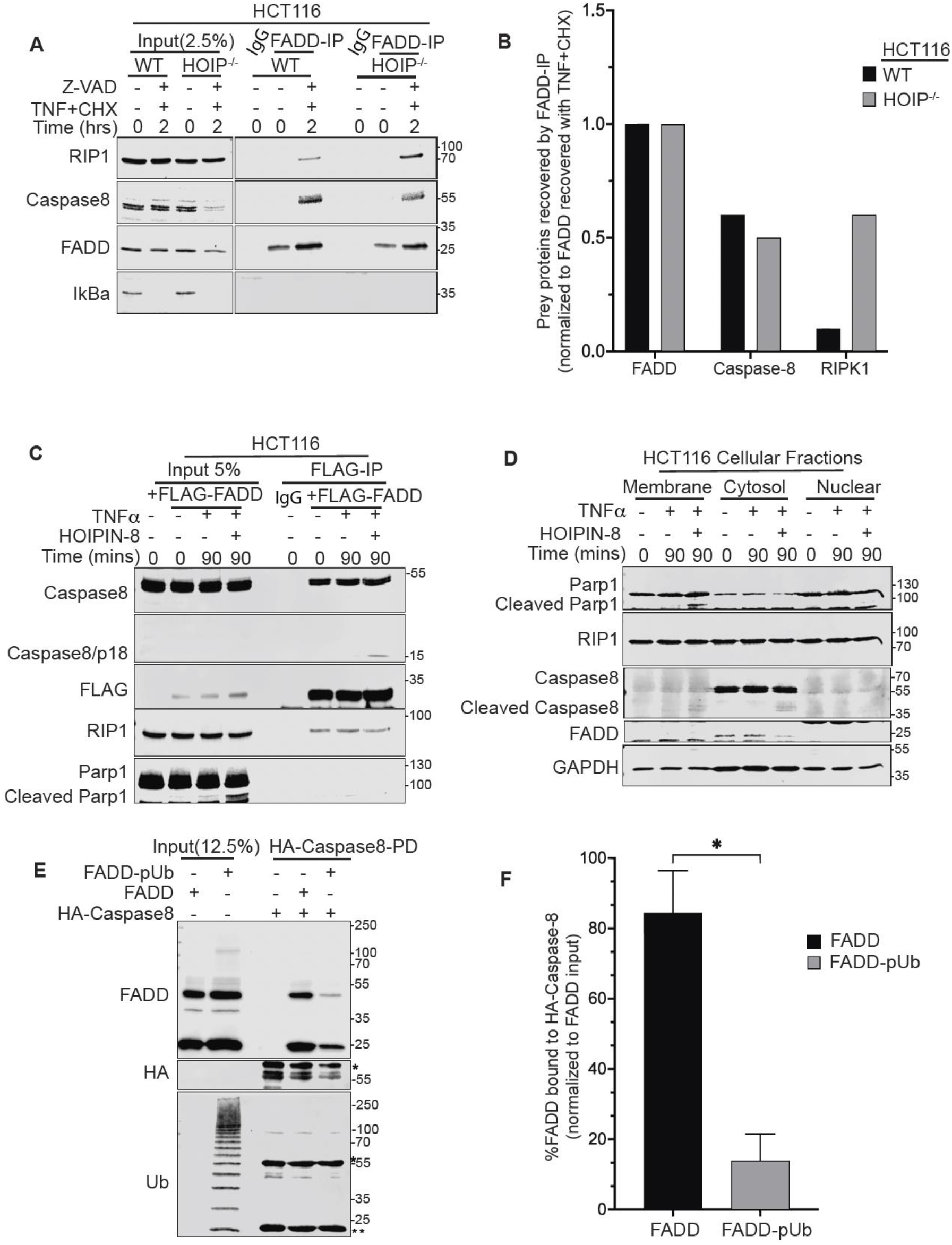
Ubiquitination of FADD reduces Caspase-8 binding, and LUBAC inhibition promotes Caspase-8 processing within complex II. **(A)** Immunoblots of endogenous FADD immunoprecipitation (IP) from WT and HOIP-KO HCT116 cells left untreated or treated with TNF [20 ng/mL] plus CHX [20 µg/mL] for 2 h, in the presence of Z-VAD-fmk (20 µM). IκBα loss in input confirms TNF signaling. Input, 2.5%; IgG, control IP. **(B)** Quantification of FADD, Caspase-8 and RIPK1 recovered in FADD IPs from (A), normalized to input. Blots were quantified by western blot densitometry using ImageJ. **(C)** FLAG IP from HCT116 cells expressing FLAG-FADD, treated with TNF [20ng/mL] with or without HOIPIN-8 (50 µM) for the indicated times. Input, 5%; IgG, control IP from untransfected cells. HOIPIN-8 increases Caspase-8 processing (p18, IP; cleaved PARP1, input) without altering the amounts of Caspase-8 or RIPK1 recovered. **(D)** Immunoblots of membrane, cytosolic, and nuclear fractions from HCT116 cells treated with TNF with or without HOIPIN-8 for 0-90 min. **(E)** Pulldown of HA-Caspase-8 with FADD or ubiquitinated FADD (FADD-pUb; immunoblotted for FADD, HA, and ubiquitin. Input, 12.5%. Asterisk (*), non-specific band. **(F)** Quantification of FADD recovered in (E), normalized to input. Blots were quantified by western blot densitometry using ImageJ. Blots are representative of at least two independent experiments.

We next asked whether pharmacological LUBAC inhibition produces a similar effect using a complementary system. To test this, we treated FLAG-FADD-overexpressing cells with TNF in the presence or absence of the HOIP catalytic inhibitor HOIPIN-8 ^31^. FLAG-FADD immunoprecipitation recovered comparable levels of Caspase-8 and RIPK1 across all conditions tested (untreated, TNF alone, and TNF+HOIPIN-8), indicating that FADD overexpression is sufficient to constitutively nucleate assembly of this complex regardless of LUBAC activity. Notably, however, the Caspase-8 cleavage fragment p18, indicative of productive, active-site proteolytic processing, was detected within the FADD-associated pool specifically under TNF+HOIPIN-8 treatment, and not under TNF alone (**Figure 3C**). This raises the possibility that LUBAC restrains a step subsequent to complex assembly, such as proteolytic processing of the recruited Caspase-8, though the present data cannot distinguish whether this reflects direct ubiquitination of Caspase-8, altered complex composition, or complex turnover kinetics.

Subcellular fractionation of HCT116 cells into membrane, cytosolic, and nuclear compartments showed that Caspase-8 and FADD were both predominantly recovered in the cytosolic fraction across all conditions tested, with comparatively little in the membrane or nuclear fractions. Cleaved PARP1 and processed Caspase-8 (p43/41) were detectable specifically within the cytosolic (and to a lesser extent membrane) fractions following TNF+HOIPIN-8 treatment (**Figure 3D**), consistent with the cytosolic pool of FADD and Caspase-8 as the relevant site of LUBAC-regulated apoptotic signaling. Finally, we tested the effect of FADD ubiquitination on Caspase-8 binding directly, using a purified, polyubiquitinated FADD species (FADD-pUb) in pulldown assays with HA-Caspase-8. Unmodified FADD was efficiently recovered in Caspase-8 pulldowns, whereas recovery of FADD-pUb was reduced under otherwise identical conditions, despite comparable input levels of each FADD species (**Figure 3E**). Thus, linear ubiquitination of FADD alone is sufficient to impair Caspase-8 binding. Together, these results indicate that FADD and Caspase-8 can assemble into a complex independently of LUBAC activity, but that LUBAC-dependent ubiquitination of FADD subsequently limits productive engagement between FADD and Caspase-8 and restrains proteolytic activation of Caspase-8 within this complex.

### Multiple death domain lysines support LUBAC-dependent FADD ubiquitination in in vitro

To determine which of FADD’s lysines are functionally capable of supporting LUBAC-dependent ubiquitination, we generated a lysine-deficient FADD background (K0) in which all eight lysines (K24, K33, K35, K110, K120, K125, K149, and K153) were substituted with arginine and then restored single lysines individually onto this lysine-less (K0) background to test each one’s sufficiency in isolation (**Figure 4A**). In reconstituted in vitro ubiquitination assays, the K0 mutant was essentially unmodified, confirming that lysines, rather than serines, account for the great majority of LUBAC-dependent FADD ubiquitination. Individual restoration of K110, K120, K149, or K153 each restored modification, whereas restoring K24, K33, or K35 individually did not (**Figure 4B**), indicating that FADD possesses multiple, partially redundant acceptor lysines concentrated within its death domain (K110–K153) rather than its death effector domain (K24–K35). Interestingly, the pattern of modification was not uniform across these four permissive sites. K120 alone supported the single most intense modified species on the blot, but this was predominantly a discrete, lower-order shift rather than an extended high-molecular-weight ladder (**Figure 4B, lane 8**). By contrast, K110, K149, and K153 each supported both a discrete shifted species and modest higher-molecular-weight smearing.

**Figure 4.**
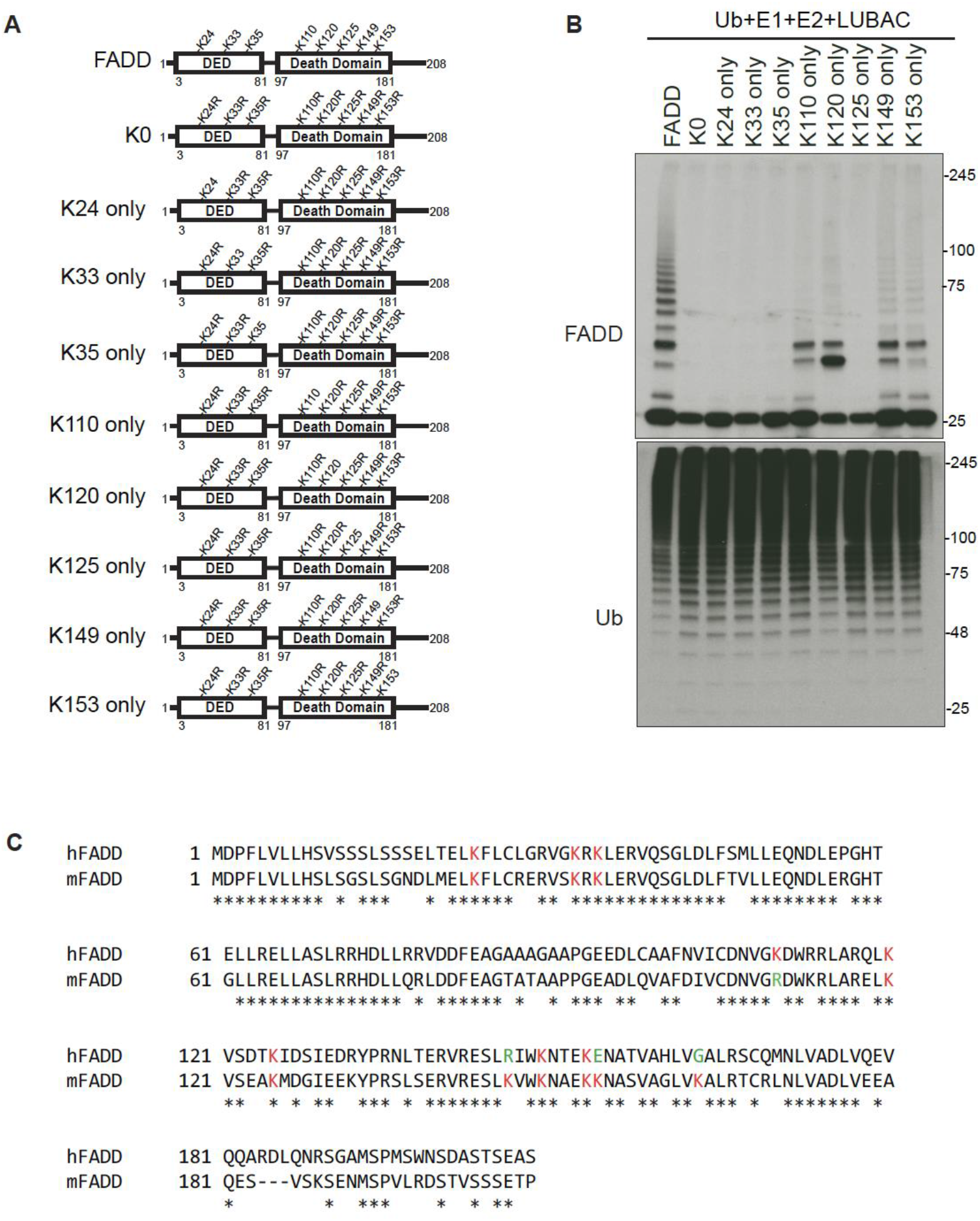
LUBAC ubiquitinates multiple lysine residues of FADD in vitro. **(A)** Schematic of FADD (208 residues; DED, residues 3-81; death domain, residues 97-181) and of the lysine-to-arginine variants used. FADD has eight lysines (K24, K33, K35, K110, K120, K125, K149, K153). K0, all eight lysines mutated to arginine; “X only,” the indicated lysine retained and the other seven mutated. **(B)** In vitro ubiquitination of WT FADD and the variants in (A) with ubiquitin, E1, E2, and LUBAC (HEK293T), immunoblotted for FADD and ubiquitin. K110, K120, K149, and K153 each support limited modification of FADD, whereas none of the single-lysine variants reproduce the WT ladder and K0 is not modified. **(C)** Alignment of human (h) and mouse (m) FADD. Lysines are highlighted in red, and green letters indicate positions where lysine is present in the corresponding sequence of the other species, and asterisks (*) mark identical residues.

To ask whether these acceptor lysines are evolutionarily conserved, and therefore of functional importance rather than incidental sites of in vitro modification, we aligned human and mouse FADD sequences. Of the four lysines capable of independently supporting robust modification, three lysines (K120, K149, and K153) are conserved in mouse FADD. K110, by contrast, is replaced by arginine in mouse, a substitution that preserves the residue’s basic charge but eliminates its capacity to serve as a ubiquitin acceptor (**Figure 4C**). This raises the possibility that K110-dependent modification is human-specific, or that its contribution is redundant with the three conserved sites in species lacking it.

### FADD and Caspase-8 are required for TNF+CHX-induced apoptosis, which is attenuated by a FADD-ubiquitin fusion

To define the contribution of FADD and Caspase-8 to TNF-induced apoptosis, we generated FADD-KO and Caspase-8-KO HCT116 cells (**Figure 5A and 5B**). Upon TNF/CHX treatment, wild-type cells showed cleavage of Caspase-8 (p43/41 and p18), Caspase-3, PARP1, and RIPK1 at 2.5 hours, and this was markedly increased in HOIP-KO cells, consistent with LUBAC restraining apoptotic signaling (**Figure 5C**). In contrast, FADD-KO and Caspase-8-KO cells showed no detectable cleavage of any of these markers, indicating that both proteins are required for apoptosis in this setting. IκBα was lost in all lines by 2.5 hours, indicating that TNFR1 signaling and CHX activity were intact. Endogenous FADD protein levels declined after TNF/CHX treatment in WT cells, more markedly in HOIP-KO cells, and to a lesser extent in Caspase-8-KO cells (**Figure 5C**). Because some loss occurred without Caspase-8, FADD turnover under these conditions is not solely a consequence of Caspase-8 activity.

**Figure 5.**
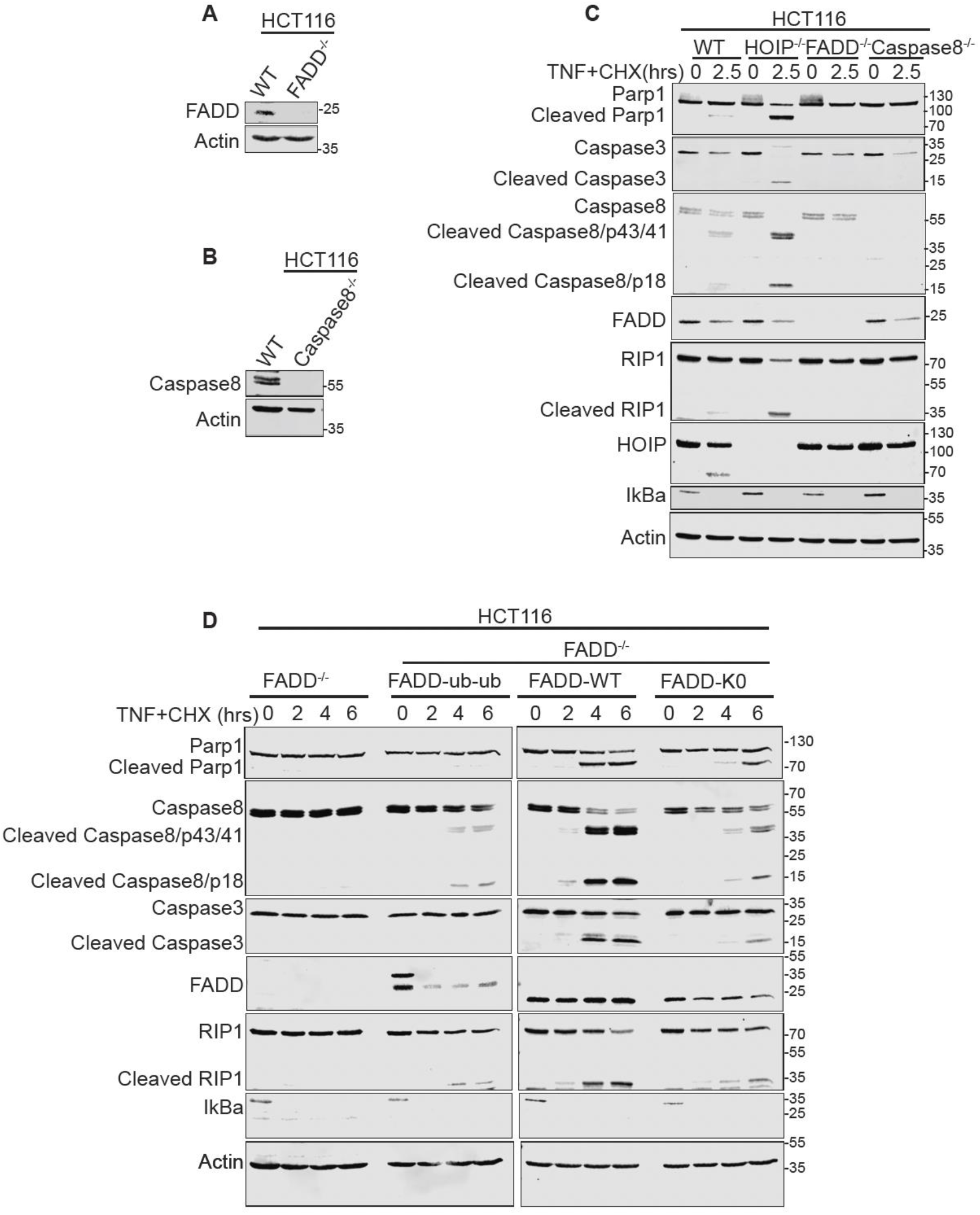
FADD and Caspase-8 are required for TNF/CHX-induced apoptosis, which is attenuated by FADD-ubiquitin fusion. **(A,B)** Immunoblots confirming loss of FADD (A) or Caspase-8 (B) in CRISPR/Cas9-generated clones of FADD-KO and Caspase-8-KO HCT116 cells. Actin, loading control. **(C)** Immunoblots of WT, HOIP-KO, FADD-KO and Caspase-8-KO HCT116 cells treated with TNF [20 ng/mL] plus CHX [20 µg/mL] for 0 or 2.5 h, probed for cleaved PARP1, Caspase-3, Caspase-8 and RIPK1 (apoptosis), IκBα (TNF signaling), and HOIP, FADD and Caspase-8 (genotype). Actin, loading control. **(D)** Immunoblots of FADD-KO HCT116 cells and FADD-KO cells stably expressing FADD-Ub-Ub (linear di-ubiquitin fused to the C-terminus of FADD), FADD-WT or FADD-K0 (all eight lysines mutated to arginine), treated with TNF [20 ng/mL] plus CHX [20 µg/mL] for 0-6 h. Dividers indicate samples run on separate gels. Blots are representative of at least two independent experiments.

**Figure 6.**
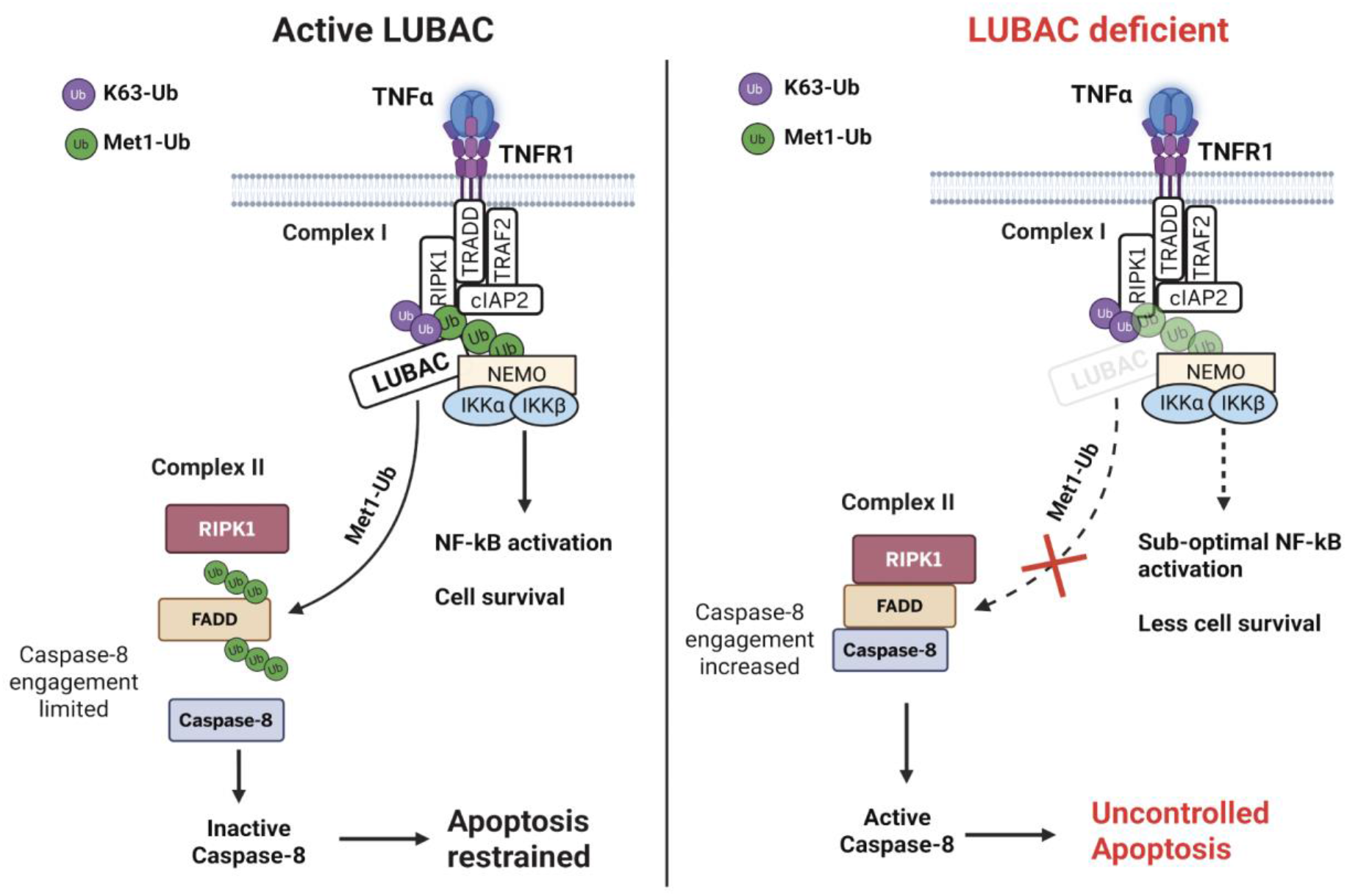
Model for LUBAC-dependent restraint of TNF-induced apoptosis at complex I and complex II. ***Left***, active LUBAC. Following TNF binding to TNFR1, LUBAC generates Met1-linked (linear) ubiquitin chains (green) within complex I, alongside K63-linked chains (purple). This stabilizes complex I and promotes NEMO-IKK-dependent NF-κB activation and cell survival. LUBAC also catalyzes Met1-linked ubiquitination of FADD in complex II, which limits FADD-Caspase-8 engagement, restricts Caspase-8 processing and restrains apoptosis. ***Right***, LUBAC deficiency. Loss of LUBAC activity reduces Met1-linked ubiquitination in complex I, resulting in sub-optimal NF-κB activation and diminished survival signaling. FADD is no longer ubiquitinated, so FADD-Caspase-8 engagement increases, Caspase-8 is activated, and cells undergo enhanced apoptosis. Dashed arrows indicate reduced or lost activity.

Reconstitution of FADD-KO cells with WT FADD restored cleavage of Caspase-8, Caspase-3, PARP1, and RIPK1, which was detectable by 2 hours, strong by 4 hours, and further increased at 6 hours (**Figure 5D**). The lysine-less mutant FADD-K0 also supported apoptotic signaling, although cleavage was delayed and reduced relative to WT. FADD-K0 levels declined over the time course, as did endogenous FADD (**Figure 5C**). In cells expressing the FADD-2Ub fusion, which mimics constitutive linear ubiquitination, cleavage of Caspase-8, Caspase-3, PARP1, and RIPK1 was markedly attenuated compared with WT FADD and was lower than in FADD-K0 cells, even though proximal TNFR1 signaling, as measured by IκBα loss, was intact. Together, these data show that TNF/CHX-induced apoptosis in HCT116 cells proceeds through FADD and Caspase-8, that LUBAC limits this pathway, and that a FADD protein carrying linear ubiquitin is impaired in promoting Caspase-8 activation. These findings are consistent with a model in which linear ubiquitination of FADD restricts assembly of a productive death-inducing complex.

## Discussion

In this study, we show that FADD and Caspase-8 are direct substrates of LUBAC that are modified through distinct routes. FADD ubiquitination requires the sequential activities of HOIL-1L and HOIP, consistent with a priming-and-extension, whereas Caspase-8 is ubiquitinated by HOIP-containing subcomplexes that lack HOIL-1L. Linear ubiquitination of FADD reduces its binding to Caspase-8 in vitro, and LUBAC activity is associated with reduced Caspase-8 processing within complex II.

A central question is what determines whether a LUBAC substrate requires HOIL-1L priming. The priming-and-extension mechanism we observe for FADD mirrors that described for LUBAC’s own subunits and for NEMO, in which HOIL-1L deposits an initial ubiquitin that HOIP subsequently extends^21,27^. Our finding that Caspase-8 bypasses this requirement indicates that this two-step mechanism is substrate-specific rather than universal. One possibility is that HOIL-1L priming is required when a substrate lacks an intrinsic feature such as a favorably positioned lysine, or a docking surface for HOIP, that allows direct engagement by HOIP alone, and that Caspase-8 possesses such a feature while FADD does not. Consistent with this, ubiquitin conjugation onto FADD promoted recruitment of HOIP specifically, suggesting that HOIL-1L priming generates a docking site that Caspase-8 may provide intrinsically. Distinguishing these will require structural characterization of how HOIP engages each substrate.

These findings extend work showing that LUBAC restrains complex II formation and signaling^18–20^, and suggests that this restraint operates at two levels – stabilization of complex I upstream^11–15^, and direct modification of complex II components once assembled, providing a second checkpoint on the apoptotic machinery. Such a model fits the reversibility of linear ubiquitination, as OTULIN and CYLD have been implicated in removing linear chains to permit death signaling^19,28^. Our finding that active but not catalytically dead OTULIN reverses FADD ubiquitination in vitro supports dynamic regulation, in which chain removal may be a decisive step toward apoptosis.

These findings may also provide a mechanistic framework for a genetic observation in LUBAC-deficient tissue. In Sharpin-deficient mice, TRADD-dependent TNFR1 signaling in epidermal keratinocytes drives skin inflammation that is prevented by epidermal FADD deletion combined with RIPK3 deficiency, and FADD or TRADD depletion suppresses the TNF sensitivity of Sharpin-deficient HaCaT cells^32^. That work established FADD as a critical effector of LUBAC-restrained apoptosis but not how LUBAC acts on it. Our data suggest the ubiquitination of FADD and Caspase-8 may directly limit their engagement within complex II. This link is inferential, as we did not test whether FADD ubiquitination is altered by Sharpin loss in keratinocytes.

FADD is modified on multiple, partially redundant death domain lysines, a configuration that would make the regulatory switch robust to loss of any single site. Three of the four validated sites are conserved in mouse, whereas K110 appears human-specific, suggesting possible species-specific tuning that warrants examination in primary or animal-derived cells.

Several limitations of this study should be noted. Reconstituted reactions with recombinant proteins establish direct, substrate-intrinsic requirements but may not reflect the kinetics, stoichiometry and regulation of LUBAC in cells. Ubiquitin-fusion (FADD-Ub, and Caspase-8-Ub) are static and may not reproduce the chain length or topology generated by LUBAC in cells, or the reversibility suggested by our OTULIN experiments. Functional experiments were restricted to HCT116 cells and TNF. Extension to keratinocytes, where genetic evidence implicates FADD-dependent apoptosis in LUBAC-deficient cells, to TRAIL and FasL, and in vivo settings is needed. Whether the priming-independent mechanism we describe for Caspase-8 extends to RIPK1 or RIPK3 also remains an open.

In summary, FADD and Caspase-8 as direct LUBAC substrates ubiquitinated through distinct catalytic mechanisms, and ubiquitination of FADD limits productive FADD-Caspase-8 engagement. These findings add a substrate-specific dimension to our understanding of how a single ubiquitin ligase complex may achieve selective control over the decision between survival and death downstream of TNF.

## Materials and Methods

### Cell lines and culture conditions

The human colorectal carcinoma cell line HCT116, used as the primary experimental model, as well as the human embryonic kidney cell line HEK293T were obtained from ATCC. Cell lines were maintained in DMEM, 1X (Dulbecco’s Modified of Eagle’s Medium with 4.5 g/L glucose, L-glutamine, and sodium pyruvate) (Corning), supplemented with 1% penicillin-streptomycin (Cytiva) and 10% FBS (Corning). Cell lines were cultured at 37°C in a humidified environment with 5% Carbon Dioxide. Cell lines were routinely tested for Mycoplasma contamination.

### CRISPR/Cas9 KO cells generation

Knockout (KO) HCT116 cells for HOIP, FADD, and Caspase-8 were generated using the CRISPR/Cas9 technique. Briefly, two guide RNAs for each gene (Supplementary Table 1) were cloned into the pSpCas9(BB)-2A-Puro (pSpCas9(BB)-2A-Puro (PX459) was a gift from Feng Zhang Addgene plasmid # 48139 ; http://n2t.net/addgene:48139 ; RRID:Addgene_48139) by restriction enzyme cloning and verified by plasmid sequencing. HCT116 cells in DMEM supplemented with 2% FBS were transfected with guide RNAs targeting each gene using Lipofectamine 3000 (Thermo Fisher Scientific) according to the manufacturer’s instructions. Following 24 hours post-transfection, the medium was replaced with complete DMEM supplemented with 0.75 µg/mL Puromycin) and the cells were cultured for 2 to 3 days. Cells were allowed to grow out, and limiting dilution to single cells in a 96-well plate was performed to obtain clones. HOIP, FADD, and Caspase-8 knockouts were verified by Western blotting.

### Constructs and stable cell lines

Human FADD constructs, FADD WT, FADD-K0 and FADD-Ub-Ub were generated and cloned into pRetro-x-tight retroviral vector (Takara). Retroviral particles for these constructs were generated in HEK293T cells by co-transfecting the constructs with the viral packaging plasmids, pCAG-VSVG (pCAG-VSVG was a gift from Arthur Nienhuis & Patrick Salmon Addgene plasmid # 35616 ; http://n2t.net/addgene:35616 ; RRID:Addgene_35616) and gag/Pol (gag/pol was a gift from Tannishtha Reya Addgene plasmid # 14887 ; http://n2t.net/addgene:14887 ; RRID:Addgene_14887), using TurboFect (Thermo Fisher Scientific) according to the manufacturer’s protocol. FADD knockout HCT116 cells were stably reconstituted with FADD WT, FADD-K0, and FADD-Ub-Ub, were generated by transduction with the respective retroviral particles. After 48 hours post-transduction, knockout cells re-expressing the constructs were selected with 2.5 µg/mL Puromycin for 5 to 10 days and verified by Western blotting.

### Recombinant protein expression and purification

Human FADD WT, FADD-K0, FADD-Ub, OTULIN WT, OTULIN C129S, UbcH5b, and mouse NEMO-UBAN were cloned into the pGEX-6P-1 vector with an N-terminal GST tag and expressed in *E. coli* BL21(DE3) in LB medium using 0.3 mM IPTG induction for 16 hours at 25°C. Cells were harvested, lysed in either 1x PBS pH 7.4 with 500 mM NaCl, 1 mM DTT, 1X protease inhibitor cocktail, and 1 mg/mL Lysozyme or buffer containing 50 mM Tris-HCl, pH 8.0, 200 mM NaCl, 2 mM EDTA, 1 mM DTT, and 1X protease inhibitor cocktail (Roche). Recombinant proteins were then affinity-purified on glutathione sepharose (Cytiva). GST-fusion proteins were cleaved from the affinity tag with 3C-protease (Acron Biosystems) in 50 mM Tris-HCl, pH 7.5, 150 mM NaCl, and 1 mM DTT. All 6xHis-tagged proteins, including UBE1 (pET28-mE1 was a gift from Jorge Eduardo Azevedo Addgene plasmid # 32534 ; http://n2t.net/addgene:32534 ; RRID:Addgene_32534), UbcH5c, UbcH7, USP2 (USP2 was a gift from Cheryl Arrowsmith Addgene plasmid # 36894 ; http://n2t.net/addgene:36894 ; RRID:Addgene_36894), OTUB1 (pOPINB-OTUB1* was a gift from David Komander Addgene plasmid # 65441 ; http://n2t.net/addgene:65441 ; RRID:Addgene_65441), and AMSH (pOPINB-AMSH* was a gift from David Komander Addgene plasmid # 66712 ; http://n2t.net/addgene:66712 ; RRID:Addgene_66712), were expressed under the same conditions, harvested, and lysed in buffer containing 50 mM Tris-HCl, pH 7.5, 200 mM NaCl, 10 mM imidazole, 5 mM β-mercaptoethanol, and 1X protease inhibitor cocktail, then purified on Ni-NTA agarose (Qiagen). Proteins were eluted with 250 mM imidazole and dialyzed overnight. To obtain the trimeric protein complex, C-terminal FLAG-tagged human HOIP, HOIL-1L, and SHARPIN were co-expressed in HEK293T cells, and lysed in Buffer H 0.3 M KCl supplemented with 1X phosphatase inhibitor and 1X protease inhibitor cocktails. The recombinant LUBAC was purified using anti-FLAG M2 beads. C-terminal FLAG-tagged NEMO and catalytically inactive Caspase-8 C360A, tagged at the N and C-terminus with FLAG and HA, respectively, were also expressed and purified from HEK293T cells under the same conditions above. The proteins were eluted in Buffer H 0.1 M KCl supplemented with 3X-FLAG peptide. The concentrations of all purified proteins were determined on NanoDrop (Thermo Fisher Scientific) using the molecular weight and extinction coefficients of each protein.

### Western blotting

Cells were lysed in lysis buffer (50 mM Tris-HCl pH 7.5, 150 mM NaCl, 1 mM EDTA, 1% Nonidet P-40) or RIPA lysis buffer (50 mM Tris-HCl pH 7.5, 150 mM NaCl, 1 mM EDTA, 0.5% sodium deoxycholate, 0.1% SDS,1% Nonidet P-40) supplemented with EDTA-free 1X protease inhibitor cocktail and 1X phosphatase inhibitor cocktail on ice for 20 mins. Protein concentrations (where indicated) were determined by BCA protein assay (Pierce^TM^). Equal amounts of protein [30 mg] were denatured in 4X Laemmli sample buffer and boiled at 95°C for 5 minutes. Harvested cells are directly lysed in 1X Laemmli sample buffer and boiled at 95°C for 5 minutes. Proteins were separated by SDS-PAGE and transferred to a nitrocellulose or PVDF membrane. Membranes were blocked in 5% milk and incubated with primary antibodies at optimal dilutions (Supplementary Table) against specific targets, followed by HRP-conjugated secondary antibodies.

### Pull-Down assay

The interaction of LUBAC with FADD or monoubiquitinated FADD was assessed by enriching for LUBAC subunits using GST (negative control), GST-FADD, or GST-tagged monoubiquitinated FADD. Briefly, recombinant FLAG-tagged Caspase-8 (positive control), HOIP, HOIL, and SHARPIN were transiently expressed separately in HEK293T cells and lysed in 1% NP-40 buffer (50 mM Tris-HCl pH 7.5, 150 mM NaCl, 1 mM EDTA, 1% NP-40 supplemented with an EDTA-free 1X protease inhibitor cocktail and a 1X phosphatase inhibitor cocktail). Each lysate was incubated with 1% NP-40-equilibrated GST, GST-FADD, or GST-FADD-Ub beads for 2 hours at 4°C with end-to-end mixing. Beads were washed 4-6 times with lysis buffer and harvested at 3000 rpm for 3 mins at 4°C. Proteins were eluted from the beads by boiling in 2x Laemmli sample buffer at 95°C for 5 mins. Proteins were then resolved on 10% SDS-PAGE gel and analyzed by western blotting.

### TNF stimulation and cell death induction

Cells designated for TNF stimulation were seeded at 2 × 10^6^ cells/mL in complete DMEM overnight prior to treatment. Cells were stimulated by replacing the media with fresh DMEM supplemented with 20 ng/mL TNF for the indicated times in each experiment. For cell death induction, cells were treated with 20 ng/mL TNF together with either 20 µg/mL cycloheximide (CHX) or 50 µM HOIPIN-8 inhibitors, with or without 20 µM zVAD-fmk, for the indicated times. When HOIPIN-8 was used to induce cell death, cells were pretreated for 1 hour prior to TNF stimulation. Proteins were separated on either 10% SDS-PAGE or a 6–20% SDS-PAGE gradient gel and analyzed by western blotting using antibodies against cell death markers.

### FADD immunoprecipitation (FADD-IP)

FADD was immunoprecipitated from both HCT116 wild-type and HOIP KO cell lines. Briefly, overnight-seeded cells were cultured in the presence of 20 ng/mL TNF, 20 µg/mL cycloheximide (CHX), and 20 µM zVAD.fmk for the indicated times, while control cells were left untreated. Cells were lysed in IP lysis buffer (30 mM Tris-HCl, pH 7.5; 120 mM NaCl; 2 mM KCl; 2 mM EDTA, pH 8; 1% Triton X-100) supplemented with an EDTA-free 1X protease inhibitor cocktail and a 1X phosphatase inhibitor cocktail. An aliquot of each treatment was saved for subsequent western blotting. Lysates were incubated with or without 1 µg/mL rabbit IgG isotype (Thermo Fisher Scientific) and anti-FADD (ProteinTech) overnight at 4°C with end-to-end mixing. Lysates incubated with the respective antibodies were subjected to IP using Protein-A Sepharose (Santa Cruz) for 4 hours at 4°C with end-to-end mixing. For Caspase-8 IP, mouse IgG isotype (Thermo Fisher Scientific), anti-Caspase-8 antibody (Cell Signaling), and Protein-G Sepharose (Santa Cruz) were used. Beads were harvested and washed 4 to 6 times with IP lysis buffer at 3000 rpm for 3 min at 4°C. Proteins were eluted from the beads and denatured by boiling in 2x Laemmli sample buffer at 95°C for 5 mins. Proteins were then resolved on a 10% SDS-PAGE gel and analyzed by western blotting.

### Denaturing immunoprecipitation

Endogenous FADD or Caspase-8 was subjected to denaturing IP to distinguish direct ubiquitination from protein association. Briefly, HCT116 cells were seeded overnight before treatment. Cells were treated with or without TNF and with TNF plus cycloheximide for the indicated times. Control cells were left untreated. Cells were lysed in RIPA lysis buffer (50 mM Tris-HCl pH 7.5, 150 mM NaCl, 1 mM EDTA, 0.5% sodium deoxycholate, 0.1% SDS, 1% NP-40) supplemented with EDTA-free 1X protease inhibitor cocktail and 1X phosphatase inhibitor cocktail. An aliquot of clarified lysate for each treatment was saved as input. For FADD-IP, lysates were incubated with or without rabbit IgG isotype and anti-FADD antibody, whereas for Caspase-8-IP, mouse IgG and Caspase-8 antibody were used. Lysates and antibodies were incubated overnight with end-to-end mixing at 4°C, then incubated with Protein-A and Protein-G Sepharose for FADD and Caspase-8, respectively, for 4 hours at 4°C with end-to-end mixing. Beads were harvested and washed 4 to 6 times at 3000 rpm for 3 minutes at 4°C. Proteins were eluted from the beads and denatured by boiling in 2x Laemmli sample buffer at 95°C for 5 minutes. Proteins were then separated on a 10% SDS-PAGE gel and analyzed by western blotting.

### Complex II isolation by FLAG-FADD immunoprecipitation

HCT116 cells were seeded and transiently transfected with 1 µg of FLAG-FADD plasmid for 16 hours prior to treatments. Transfected cells were treated with or without TNF and TNF plus HOIPIN-8 for the indicated concentrations and times. Non-transfected and transfected control cells were left untreated. Cells were lysed in FLAG-IP lysis buffer, and aliquots were saved for inputs. To isolate FLAG-FADD together with complex II components, lysates were incubated with FLAG-M2 beads for 3 hours at 4°C with end-to-end mixing. Beads were harvested and washed 4 to 6 times at 3000 rpm for 3 minutes at 4°C. Proteins were eluted from the beads and denatured by boiling in 2X Laemmli sample buffer at 95°C for 5 minutes. Proteins were then separated on a 10% SDS-PAGE gel and analyzed by western blotting.

### Cellular fractionation

HCT116 cells were seeded overnight, as before, prior to treatment. Cells were pre-treated with or without HOIPIN-8 at the indicated concentration for 1 hour and then stimulated with or without TNF at the indicated concentrations for 90 minutes. After treatment, cells were washed twice with cold PBS, harvested by scraping in PBS, and centrifuged at 4°C and 250× g for 5 minutes. The cell pellet was then resuspended in a homogenization buffer (20 mM Tris-HCl, pH 7.5; 250 mM Sucrose; 3 mM Imidazole, pH 8; 1 mM EDTA, pH 8; supplemented with an EDTA-free 1X protease inhibitor cocktail and a 1X phosphatase inhibitor cocktail) and incubated on ice for 20 minutes. Cells were mechanically homogenized by passing through a 27G needle 15 – 18 times. The homogenate was centrifuged to separate the post-nuclear fraction or supernatant (PNS) from the nuclear fraction (NF, or pellet). The pellet was washed twice with homogenization buffer for 10 minutes at 1 000 x g at 4°C and resuspended in 1x Laemmli sample buffer for subsequent western blotting. The PNS fraction was centrifuged for 1 hour at 20 000 x g at 4°C to separate the cytosolic fraction (CF) from the membrane fraction (MF). The MF was washed twice with homogenization buffer for 5 minutes at 20 000 x g at 4°C and resuspended in 1x Laemmli sample buffer. For the CF, 4X Laemmli sample buffer was added to an aliquot, and all fractions were denatured by boiling at 95°C for 5 min. Fractions were resolved on 10% SDS-PAGE gel and analyzed by western blotting.

### *In vitro* ubiquitination assay

*In vitro* ubiquitination assays were conducted in 25 µL reactions containing 20 mM Tris-HCl (pH 7.5), 5 mM MgCl2, 2 mM ATP, and 1 mM DTT, with 100 nM E1 (human UBE1 purified in-house), 1 µM E2 (commercially obtained or purified in-house, where indicated), where indicated, 50 µM ubiquitin (R&D systems) or His-tagged ubiquitin, 1 µg of recombinantly purified LUBAC (or 1 µg of each subunit of LUBAC), and 1 µg of FADD ( or other substrates, where indicated). For in vitro ubiquitination of Caspase-8, purified, HA-tagged Caspase-8, bound to magnetic HA beads or bacterially purified Caspase-8, was used as the substrate. Reactions were incubated at 37°C for at least 2 hours unless otherwise specified and stopped with 4X Laemmli sample buffer containing β-mercaptoethanol. Proteins were denatured by boiling at 95°C for 5 minutes. Control reactions, with or without substrate or LUBAC, were included to confirm reaction specificity. Proteins were resolved on a 6–20% SDS–PAGE gradient gel and analyzed by western blotting.

### FADD and polyubiquitinated FADD pull-down with Caspase-8

Linear polyubiquitinated FADD (FADD-pUb) was generated *in vitro* as described, and the reaction was stopped with 10 mM EDTA (pH 8) for downstream pull-down. FADD only or FADD-pUb were incubated with purified, HA-tagged Caspase-8 bound to beads in PD buffer (20 mM Tris-HCl pH 7.5, 150 mM NaCl and 0.5% NP-40) overnight at 4°C with end-to-end mixing. Beads were harvested and washed 6 times with PD buffer. Proteins were eluted from the beads and denatured by boiling in 2x Laemmli sample buffer at 95°C for 5 minutes. Proteins were then separated on a 10% SDS-PAGE gel and analyzed by western blotting.

### Deubiquitinase (DUBs) Assay

*In vitro* ubiquitinated FADD was generated as previously described and incubated with either recombinant wild-type OTULIN or catalytically inactive OTULIN at a final concentration of 1 μM in deubiquitination reaction buffer (50 mM Tris-HCl (pH 7.5), 50 mM NaCl, and 5 mM DTT) for the indicated time at 37°C. Reactions were stopped by adding 2x Laemmli sample buffer, then boiling at 95°C for 5 minutes. Proteins were separated on a 10% SDS-PAGE gel and analyzed by western blotting with antibodies against FADD and ubiquitin chains.

### Cleaved-Caspase-3 Immunofluorescence

HCT116 wild-type and HOIP knockout cells were seeded onto sterile glass coverslips and cultured overnight to reach approximately 60 – 80% confluence. Cells were then left untreated or treated with TNF alone or with TNF in combination with cycloheximide for the indicated concentrations and time. After treatment, cells were washed with PBS, fixed in 4% paraformaldehyde for 20 minutes at room temperature, and permeabilized with 0.2% Triton X-100 in PBS. Non-specific binding was blocked with 5% BSA in PBST. Cells were incubated with an anti-cleaved Caspase-3 primary antibody overnight at 4°C. Cells were then washed and incubated with an appropriate fluorophore-conjugated secondary antibody, followed by nuclear counterstaining with DAPI. Coverslips were mounted with antifade mounting medium (Mowiol) and imaged by confocal microscopy (Cytation C10, Biotek) using identical acquisition settings across all experimental conditions. Cleaved Caspase-3 activation was quantified as the percentage of cleaved Caspase-3-positive cells relative to the total number of DAPI-positive nuclei from multiple randomly selected fields per condition.

### NF-kB Luciferase assay

Wild-type and HOIP KO HCT116 cells were infected with a lentivirus carrying the NF-κB p50/p65 pathway luciferase reporter plasmid with green fluorescent protein (GFP), pLminP_Luc2P_RE1 (pLminP_Luc2P_RE1 was a gift from Ramnik Xavier Addgene plasmid # 90335) . Cells stably expressing the reporter were selected by flow cytometry based on GFP intensity. For the luciferase assay, cells were seeded at 50 000 cells per 96-well black clear-bottom tissue culture plate in triplicate and incubated overnight. Cells were treated with or without TNF for 12 hours. GFP fluorescence intensities were measured after treatment using a Cytation C10 confocal imaging reader (Agilent BioTek). Relative light units were measured using a Cytation C10 confocal imaging reader (Agilent BioTek) by transferring 10 µL of cell lysates into 96-well white-bottom plates and adding 50 µL of firefly luciferase substrate (GoldBio). Luciferase activity data for wild-type and HOIP-deficient HCT116 cells were presented as fold changes after normalization to GFP fluorescence intensities.

### Data availability

Any additional information required to reanalyze the data reported in this paper is available from the corresponding author upon request. Source data are provided in this paper.

## Acknowledgements

We thank all past and current Lopez-Mosqueda Lab members.

## Contributions

Conceptualization: F.K-A., and J.L-M. Data Acquisition: F.K-A., A.B., and J.L-M. Funding Acquisition: J.L-M. Supervision: J.L-M. Writing – review and editing: F.K-A., A.B., and J.L-M.

---

**Supplemental Figure 1.**
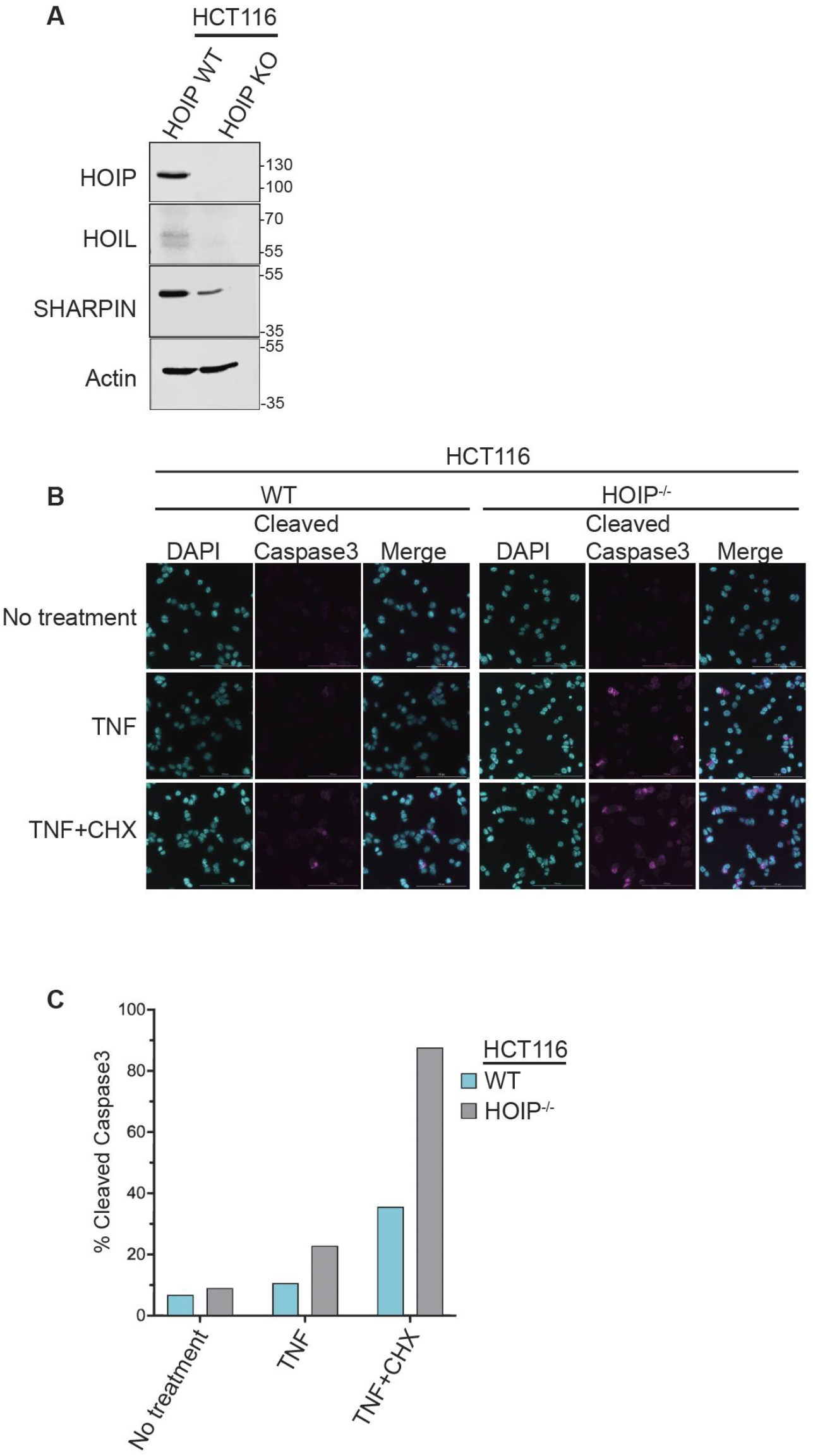
**(A)** Immunoblots confirming that loss of HOIP affects the stability of HOIL-1L and SHARPIN. **(B)** Cleaved Caspase-3 immunofluorescence staining of wild-type and HOIP knockout HCT116 cells under untreated, TNF [20 ng/mL], and TNF+CHX [20 mg/mL] treated conditions, confirming that loss of HOIP leads to increased cell death. **(C)** Quantification of the percentage of cleaved Caspase-3 IF-positive cells as a measure of cell death. Mean ± SD, n = [3].

**Supplemental Figure 2.**
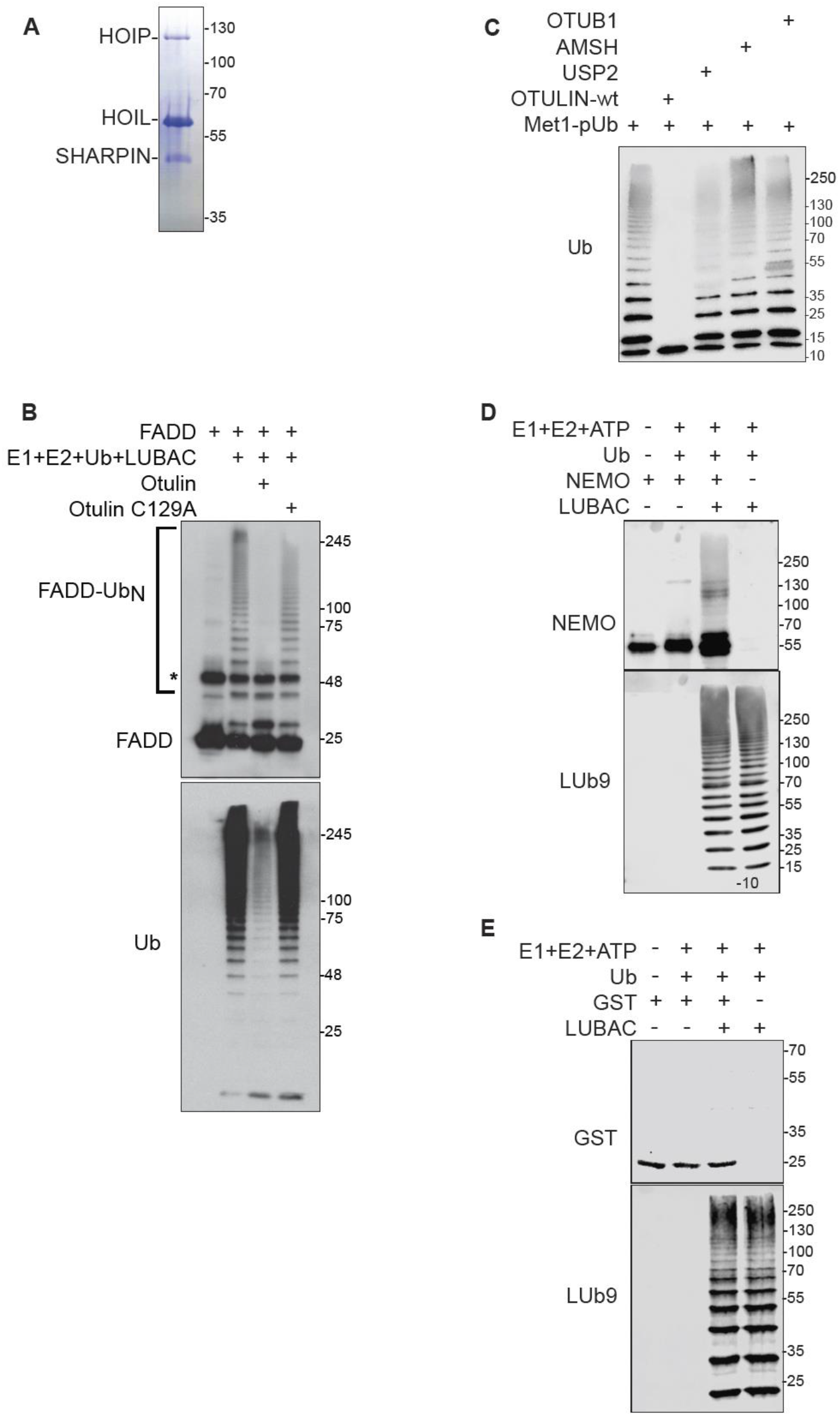
**(A)** FLAG-tag affinity purification of LUBAC after co-transfection and overexpression of FLAG-tagged HOIP, HOIL-1L, and SHARPIN in HEK293T cells. **(B)** Deubiquitinase (DUB) assay using OTULIN WT and catalytically dead OTULIN C129A on in vitro linear ubiquitinated FADD, confirming that LUBAC modifies FADD with linear ubiquitin chains. **(C)** DUB assay using OTUB1 (hydrolyzes K48 ubiquitin chains), AMSH (hydrolyzes K63 ubiquitin chains), USP2 (hydrolyzes all ubiquitin chains), and OTULIN (hydrolyzes linear ubiquitin chains) to confirm that the purified LUBAC can generate free linear ubiquitin chains. **(D)** In vitro ubiquitination of NEMO, a known substrate of LUBAC and serves as a positive control for LUBAC specificity. **(E)** In vitro ubiquitination of GST tag, as negative control for the in vitro ubiquitination assays to confirm the specificity of LUBAC in targeting its substrates.

## References

1 van Loo, G. & Bertrand, M. J. M. Death by TNF: a road to inflammation. Nat Rev Immunol 23, 289–303 (2023). 10.1038/s41577-022-00792-3

2 Micheau, O. & Tschopp, J. Induction of TNF receptor I-mediated apoptosis via two sequential signaling complexes. Cell 114, 181–190 (2003). 10.1016/s0092-8674(03)00521-x

3 Silke, J. The regulation of TNF signalling: what a tangled web we weave. Curr Opin Immunol 23, 620–626 (2011). 10.1016/j.coi.2011.08.002

4 Mahoney, D. J. et al. Both cIAP1 and cIAP2 regulate TNFalpha-mediated NF-kappaB activation. Proc Natl Acad Sci U S A 105, 11778–11783 (2008). 10.1073/pnas.0711122105

5 Shi, J. H. & Sun, S. C. Tumor Necrosis Factor Receptor-Associated Factor Regulation of Nuclear Factor kappaB and Mitogen-Activated Protein Kinase Pathways. Front Immunol 9, 1849 (2018). 10.3389/fimmu.2018.01849

6 Haas, T. L. et al. Recruitment of the linear ubiquitin chain assembly complex stabilizes the TNF-R1 signaling complex and is required for TNF-mediated gene induction. Mol Cell 36, 831–844 (2009). 10.1016/j.molcel.2009.10.013

7 Brenner, D., Blaser, H. & Mak, T. W. Regulation of tumour necrosis factor signalling: live or let die. Nat Rev Immunol 15, 362–374 (2015). 10.1038/nri3834

8 Wertz, I. E. & Dixit, V. M. Ubiquitin-mediated regulation of TNFR1 signaling. Cytokine Growth Factor Rev 19, 313–324 (2008). 10.1016/j.cytogfr.2008.04.014

9 Verhelst, K., Carpentier, I. & Beyaert, R. Regulation of TNF-induced NF-kappaB activation by different cytoplasmic ubiquitination events. Cytokine Growth Factor Rev 22, 277–286 (2011). 10.1016/j.cytogfr.2011.11.002

10 Schmukle, A. C. & Walczak, H. No one can whistle a symphony alone - how different ubiquitin linkages cooperate to orchestrate NF-kappaB activity. J Cell Sci 125, 549–559 (2012). 10.1242/jcs.091793

11 Kirisako, T. et al. A ubiquitin ligase complex assembles linear polyubiquitin chains. EMBO J 25, 4877–4887 (2006). 10.1038/sj.emboj.7601360

12 Gerlach, B. et al. Linear ubiquitination prevents inflammation and regulates immune signalling. Nature 471, 591–596 (2011). 10.1038/nature09816

13 Ikeda, F. et al. SHARPIN forms a linear ubiquitin ligase complex regulating NF-kappaB activity and apoptosis. Nature 471, 637–641 (2011). 10.1038/nature09814

14 Tokunaga, F. et al. SHARPIN is a component of the NF-kappaB-activating linear ubiquitin chain assembly complex. Nature 471, 633–636 (2011). 10.1038/nature09815

15 Tokunaga, F. et al. Involvement of linear polyubiquitylation of NEMO in NF-kappaB activation. Nat Cell Biol 11, 123–132 (2009). 10.1038/ncb1821

16 Peltzer, N. et al. LUBAC is essential for embryogenesis by preventing cell death and enabling haematopoiesis. Nature 557, 112–117 (2018). 10.1038/s41586-018-0064-8

17 Boisson, B. et al. Immunodeficiency, autoinflammation and amylopectinosis in humans with inherited HOIL-1 and LUBAC deficiency. Nat Immunol 13, 1178–1186 (2012). 10.1038/ni.2457

18 de Almagro, M. C., Goncharov, T., Newton, K. & Vucic, D. Cellular IAP proteins and LUBAC differentially regulate necrosome-associated RIP1 ubiquitination. Cell Death Dis 6, e1800 (2015). 10.1038/cddis.2015.158

19 Draber, P. et al. LUBAC-Recruited CYLD and A20 Regulate Gene Activation and Cell Death by Exerting Opposing Effects on Linear Ubiquitin in Signaling Complexes. Cell Rep 13, 2258–2272 (2015). 10.1016/j.celrep.2015.11.009

20 Lafont, E. et al. The linear ubiquitin chain assembly complex regulates TRAIL-induced gene activation and cell death. EMBO J 36, 1147–1166 (2017). 10.15252/embj.201695699

21 Goto, E. & Tokunaga, F. Decreased linear ubiquitination of NEMO and FADD on apoptosis with caspase-mediated cleavage of HOIP. Biochem Biophys Res Commun 485, 152–159 (2017). 10.1016/j.bbrc.2017.02.040

22 Carrington, P. E. et al. The structure of FADD and its mode of interaction with procaspase-8. Mol Cell 22, 599–610 (2006). 10.1016/j.molcel.2006.04.018

23 Sandu, C. et al. FADD self-association is required for stable interaction with an activated death receptor. Cell Death Differ 13, 2052–2061 (2006). 10.1038/sj.cdd.4401966

24 Riley, J. S., Malik, A., Holohan, C. & Longley, D. B. DED or alive: assembly and regulation of the death effector domain complexes. Cell Death Dis 6, e1866 (2015). 10.1038/cddis.2015.213

25 Kelsall, I. R., Zhang, J., Knebel, A., Arthur, J. S. C. & Cohen, P. The E3 ligase HOIL-1 catalyses ester bond formation between ubiquitin and components of the Myddosome in mammalian cells. Proc Natl Acad Sci U S A 116, 13293–13298 (2019). 10.1073/pnas.1905873116

26 Rodriguez Carvajal, A., et al. The linear ubiquitin chain assembly complex (LUBAC) generates heterotypic ubiquitin chains. Elife 10 (2021). 10.7554/eLife.60660

27 Fuseya, Y. et al. The HOIL-1L ligase modulates immune signalling and cell death via monoubiquitination of LUBAC. Nat Cell Biol 22, 663–673 (2020). 10.1038/s41556-020-0517-9

28 Smit, J. J. et al. The E3 ligase HOIP specifies linear ubiquitin chain assembly through its RING-IBR-RING domain and the unique LDD extension. EMBO J 31, 3833–3844 (2012). 10.1038/emboj.2012.217

29 Hahn, T. et al. Use of monoclonal antibodies to a human cytotoxin for its isolation and for examining the self-induction of resistance to this protein. Proc Natl Acad Sci U S A 82, 3814–3818 (1985). 10.1073/pnas.82.11.3814

30 Kirstein, M. & Baglioni, C. Tumor necrosis factor induces synthesis of two proteins in human fibroblasts. J Biol Chem 261, 9565–9567 (1986).

31 Katsuya, K. et al. Small-molecule inhibitors of linear ubiquitin chain assembly complex (LUBAC), HOIPINs, suppress NF-kappaB signaling. Biochem Biophys Res Commun 509, 700–706 (2019). 10.1016/j.bbrc.2018.12.164

32 Kumari, S. et al. Sharpin prevents skin inflammation by inhibiting TNFR1-induced keratinocyte apoptosis. Elife 3 (2014). 10.7554/eLife.03422

